# Comparative transcriptomic profile reveals candidate genes manipulated by type III effectors of *Pantoea agglomerans* pv. *betae* leading to gall formation in beet

**DOI:** 10.1101/2024.11.24.624952

**Authors:** Priya Gupta, Metsada Pasmanik-Chor, Hanita Zemach, Isaac Barash, Doron Teper, Guido Sessa

**Author notes:** Deceased (Guido Sessa). Correspondence to: Doron Teper, Institute of Plant Protection, Dept. of Plant Pathology and Weed Research, Agricultural Research Organization – Volcani Institute. 7505101 Rishon LeZion, Israel., Priya Gupta, School of Plant Sciences and Food Security, Tel-Aviv University, 69978 Tel-Aviv, Israel.

## Abstract

*Pantoea agglomerans* pv. *betae* (*Pab*) induces tumor-like galls in beet and gypsophila, a process mediated by the secretion of effector proteins via *Pab*’s type III secretion system (T3SS). The molecular mechanisms underlying *Pab*-induced gall formation remain largely unexplored. This study delves into the cellular architecture and transcriptional profile of *Pab*-mediated galls, comparing host responses to wild-type *Pab* and a T3SS-inactive mutant, *hrcC*^−^. Morphological analysis using scanning electron microscopy and cross-sectional visualization of infected beet leaf tissues revealed that *Pab*-induced gall-like structures are linked to cell hyperplasia and tissue ruptures, contingent on T3SS activity. Comparative transcriptome analysis of wild-type *Pab* and *hrcC*^−^ *Pab*-infected beet leaves at 12 and 48 hours unveiled significant transcriptional reprogramming, with nearly 2,000 differentially expressed genes at 48 hours post inoculation. Enrichment analyses identified the upregulation of pathways related to signal transduction, defense, carbohydrate metabolism, and cell wall modulation in wild-type *Pab*-infected leaves compared to controls. Particularly notable was the significant upregulation of numerous genes associated with cell wall loosening by wild-type *Pab*, suggesting an initial rearrangement of cell wall architecture facilitates gall formation. Furthermore, transcriptome analysis demonstrated that wild-type *Pab* suppresses the expression of the betalain biosynthetic gene *DOPA 4,5-DIOXYGENASE*, leading to reduced betalain accumulation in infected tissues compared to the mutant strain. These findings offer fresh insights into the transcriptional and physiological manipulation of host tissue during the early stages of *Pab*-induced gall formation.

## Introduction

Plant galls are a form of altered developmental trajectory, primarily associated with cell division, expansion, and altered cell identity, induced by environmental stress or various pathogens such as viruses, bacteria, fungi, nematodes, mites, and insects (Harris and Pitzschke 2020). Galls morphology and complexity vary greatly according to their associated pathogen, and they offer them advantages by serving as a microhabitat that provides sustenance, shelter, and protection against adversaries (Gatjens 2019). Many bacterial pathogens, such as *Agrobacterium tumefaciens*, *Pseudomonas savastanoi* pv. *savastanoi*, *Xanthomonas citri* pv. *citri*, and gall-forming *Pantoea agglomerans*, promote the formation of tumor-like gall structures by inducing uncontrolled cell divisions and expansion, resulting in hyperplasia and hypertrophy (Weisberg et al. 2023; Ramos et al. 2012; Ference et al. 2018; Barash and Manulis-Sasson 2009). These pathogens utilize diverse strategies to manipulate cell identity to promote gall formation, such as manipulating host transcriptional programming using pathogen effectors, altering the balance of plant hormones through manipulation of hormone production and transport, or biosynthesizing plant hormones (Harris and Pitzschke 2020). For instance, *Agrobacterium tumefaciens* promotes gall formation by translocating a T-DNA-protein complex into the plant cell through the type IV secretion system, which integrates into the plant genome and encodes genes associated with auxin, cytokinin, and opine biosynthesis, thereby promoting cell division (Weisberg et al. 2023). On the other hand, *Xanthomonas citri* pv. *citri* promotes cell division and expansion by translocating a bacterial-encoded eukaryotic transcription factor, PthA4, through the type

III secretion system (T3SS), which induces the expression of the citrus transcriptional regulator *CsLOB1* (Hu et al. 2014). This, in turn, induces the expression of genes associated with cell cycle control and cell wall loosening (Duan et al. 2018).

The *P. agglomerans* pathovars, *P. agglomerans* pv. *gypsophilae* (*Pag*) and *P. agglomerans* pv. *betae* (*Pab*), are tumorigenic pathogens, causing galls in gypsophila (*Gypsophila paniculata*) or gypsophila and beet (*Beta vulgaris*), respectively (Barash and Manulis-Sasson 2009). These pathovars have evolved from *P. agglomerans*, typically found as a commensal plant endophyte and epiphyte (Walterson and Stavrinides 2015), by acquiring a pathogenicity plasmid, designated as pPATH (Barash and Manulis-Sasson 2009; Geraffi et al. 2023). *Pag* galls in gypsophila cuttings appear as spongy undifferentiated tissue and are associated with cell expansion and cell division (Chalupowicz et al. 2006) while *Pab* galls in beet, which have yet to be anatomically characterized, appear as large undifferentiated tumors on beetroot tissues (Burr et al. 1991). The main tumor-associated determinants of *Pag* and *Pab* are encoded by the pPATH plasmid, which include an Hrp1-like T3SS, 9 (for *Pag*) or 12 (for *Pab*) T3SS effector proteins, auxin biosynthesis genes (*iaaM* and *iaaH*), and cytokinin biosynthesis genes (*etz* and *pre-etz*) (Geraffi et al. 2023). Disruption of either auxin or cytokinin biosynthesis genes in *Pag* significantly reduces gall size in gypsophila but does not eliminate gall formation (Lichter et al. 1995), while *Pag*-mediated gall formation is completely inhibited by auxin polar transport inhibitors (Chalupowicz et al. 2013), indicating that plant-synthesized hormones are essential for gall formation, while bacteria-synthesized hormones contribute but are not essential for gall formation. The T3SS is essential for gall formation in gypsophila and beet by *Pag* or *Pab*, and mutants of structural components of the T3SS are unable to cause disease symptoms (Nizan et al. 1997; Geraffi et al. 2023). Additionally, ectopic expression of the *Pab* T3SS effectors HsvB and PseB or the *Pag* T3SS effectors HsvG and PthG in non-pathogenic *P. agglomerans, Pseudomonas fluorescens*, and *Escherichia coli* that encode the T3SS apparatus is sufficient for tumor induction in gypsophila or beet (Nissan et al. 2019). While most bacterial pathogenicity determinants are well-established, the plant molecular mechanisms that facilitate gall formation by *Pag* or *Pab* are still elusive and have yet to be subjected to in-depth molecular characterization. In the study, we delineated the transcriptional response of beet leaves to *Pab*, unveiling an array of genetic signatures orchestrating the early stages of gall formation.

## Results

### *Pab* colonize and induce gall formation on beet leaves

We aimed to characterize the molecular and physiological features of *Pab*-mediated gall formation by comparing host response to wild-type *Pab* and *Pab*hrcC^−^, a mutant in an essential structural component of the type III secretion system (T3SS) that is unable to induce gall formation during root inoculations (Geraffi 2021). *Pab*-mediated gall mainly occurs in root storage tissue. However, this tissue can only be infected by surface inoculation, which result in uneven exposure to the pathogen. Therefore, we estimated whether leaf tissue, which can be directly infiltrated through a needleless syringe, could be alternatively used to assess host response. We first assessed how infiltration of *Pab* affects leaf tissue morphology and symptom development. To evaluate whether *Pab* can induce gall formation in leaf tissues, beet leaves were infiltrated with buffer control (mock), *Pab* (*Pab*WT), and *Pab*hrcC^−^ using a needleless syringe and monitored for tissue alterations (Fig. 1A and 1B). For comparison, inoculation experiments were conducted in parallel on beetroot cubes (Fig 1C). Gall formation was observed on *Pab*WT-streaked beetroot cubes at 6 days post-inoculation (dpi), whereas mock-treated and *Pab*hrcC^−^-treated cubes did not exhibit any visible disease symptoms (Fig. 1C). In agreement with this observation, *Pab*WT-infiltrated leaves show curling at 10 dpi and galls at 20 dpi, especially at the syringe-infiltrated sites, compared to mock and *Pab*hrcC^−^ inoculated leaves (Fig. 1A and B). On the other hand, *Pab*hrcC^−^ infiltrated leaves did not develop gall-like structures and showed a high level of betalain pigment accumulation compared to mock– and *Pab*WT-treated leaves (Fig. 1A and B).

**Fig. 1.**
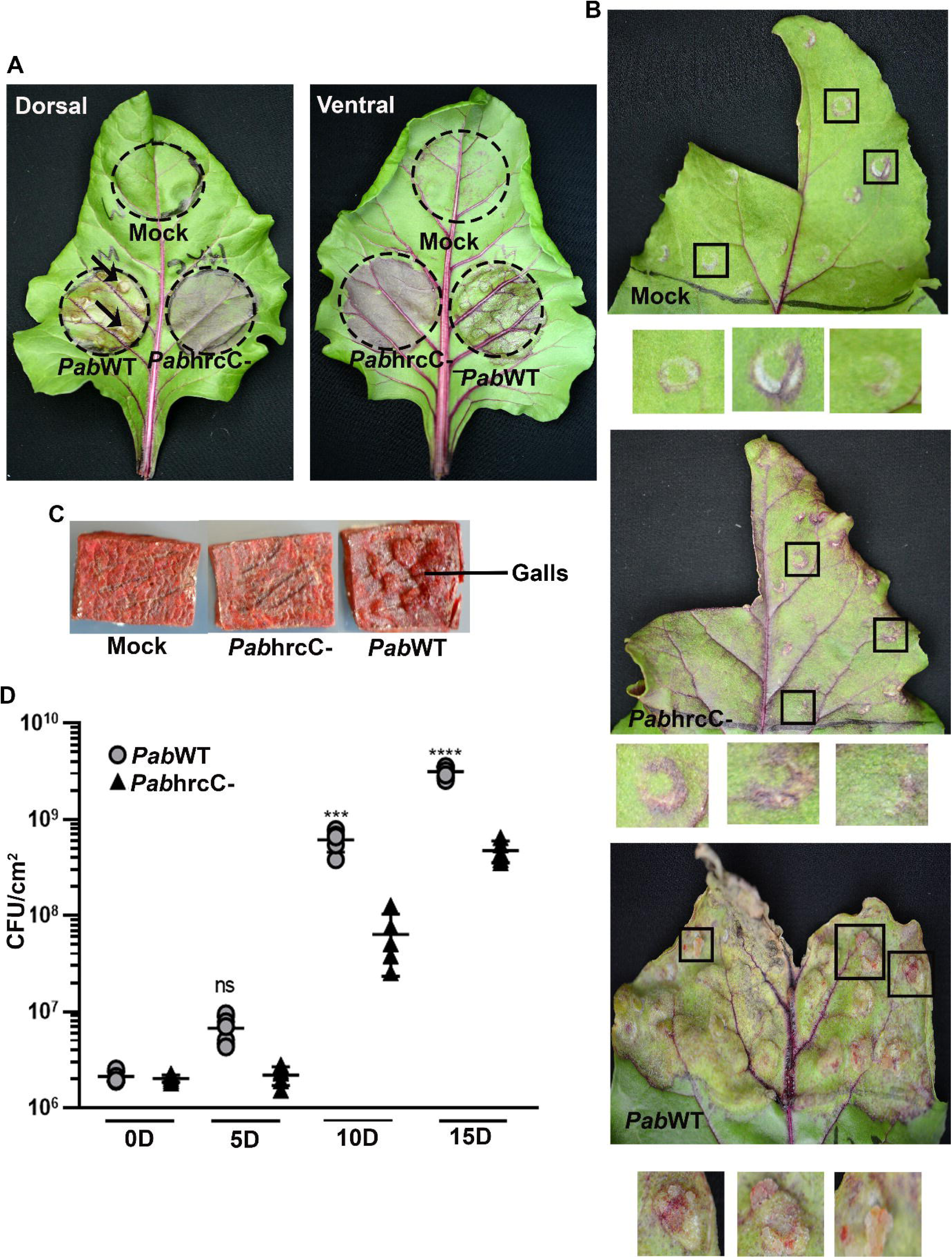
Symptoms and CFU count after mock, *Pab*WT, and *Pab*hrcC inoculations. (A, B, D) ddH2O (mock) or bacterial suspensions (10^8^ CFU/ml) of *Pab*WT or *Pab*hrcC^−^ were infiltrated into beet leaves. (A) Representative pictures depict symptoms at 10 dpi on dorsal and ventral surfaces of infiltrated leaves. (B) Representative pictures depict symptoms at 20 dpi. The insets in the figure were zoomed for better visualization of symptoms. (C) Beet root cubes were streaked with mock or 10^8^ CFU/ml of *Pab*WT or *Pab*hrcC^−^. Representative pictures were taken at 6 dpi. (D) Leaf bacterial populations were determined at 0, 5, 10, and 15 dpi. The asterisks represents a statistically significant difference in the treatments compared to *Pab*WT based on five biological replicates per treatment at each time point. It is determined by the one-way ANOVA method (Tukey’s multiple comparisons test). ns – not significant, ***(p ≤ 0.001), ****(p ≤ 0.0001). The experiments were repeated twice with similar results.

Next, we assessed the contribution of the T3SS to leaf tissue colonization. We observed a significant reduction in leaf colonization by *Pab*hrcC^−^ compared to *Pab*WT at 10 and 15 dpi (Fig. 1D); supporting that the T3SS contribute to growth in beet leaf tissue.

These findings demonstrate that *Pab* virulence on beet is not tissue-specific and it can colonize and promote the appearance of gall-like structures on beet leaf in a T3SS-dependent manner.

### *Pab* induces T3SS-dependent hyperplasia and anatomical changes in beet leaves

In order to get a deeper insight into the *Pab*-induced tissue alterations, we focused on the morphological and anatomical changes that occur in beet leaves after inoculations. Beet leaves were syringe-infiltrated with mock, *Pab*hrcC^−^, and *Pab*WT, and tissues were collected from syringe-infiltrated areas at 10 and 20 dpi and tissue alterations were visualized using scanning electron microscopy (SEM). In mock and *Pab*hrcC^−^ treated leaves, the epidermis appeared to be similarly patterned, having guard and subsidiary cells at 10 and 20 dpi (Fig. 2A and B). In contrast, the *Pab*WT-infiltrated leaves show curling and ruptured epidermis at 10 dpi, and galls with an irregular surface at 20 dpi (Fig. 2A and B).

**Fig. 2.**
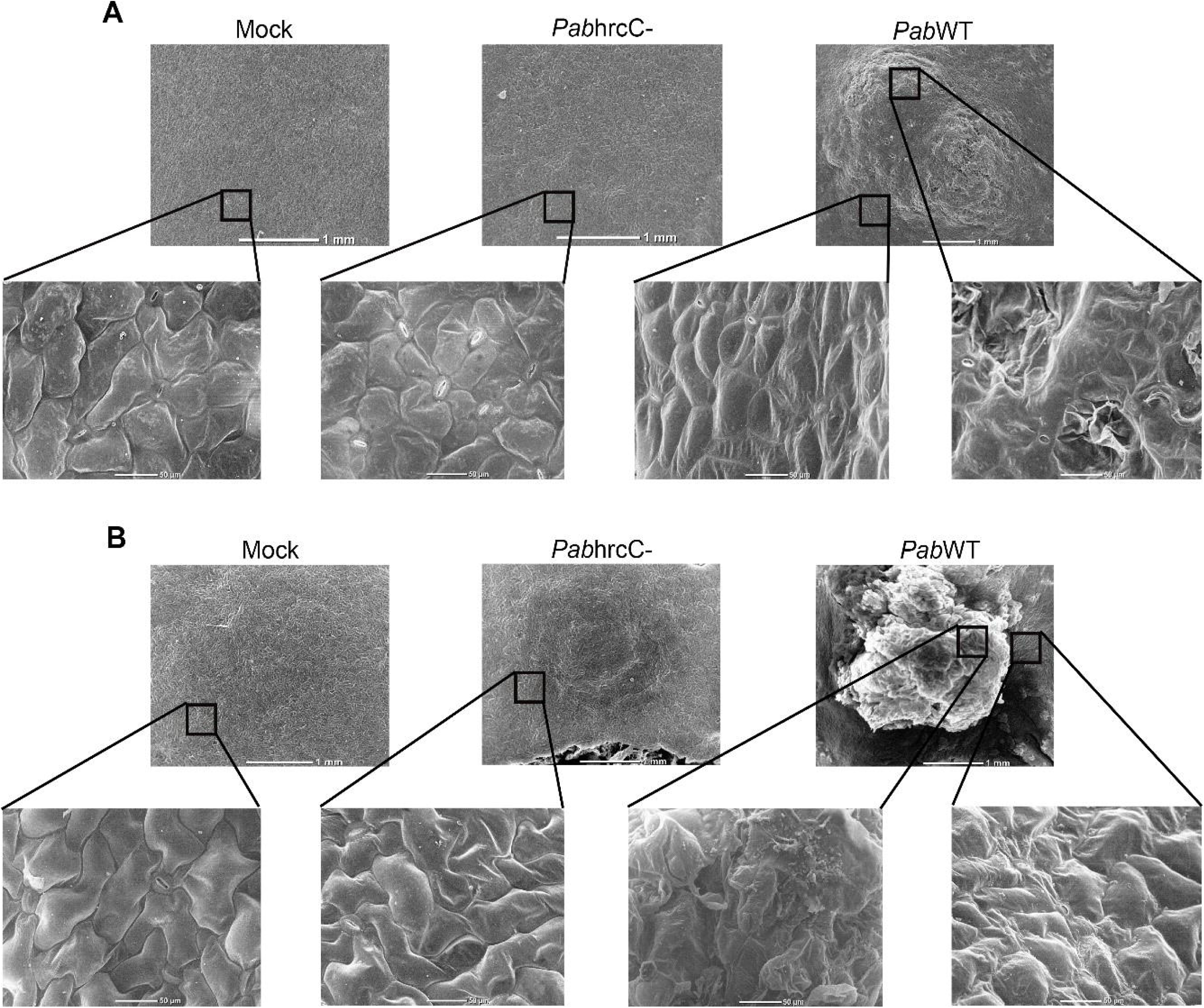
A scanning electron microscope (SEM) images of *Pab*-inoculated leaf tissues. SEM images at (A) 10 dpi (B) 20 dpi after mock (ddH_2_O), *Pab*hrcC^−^ (10^8^ CFU/ml) and *Pab*WT (10^8^ CFU/ml) infiltration in beet leaves. The inset shows the selected area for higher magnification view. The experiments were repeated twice, with five independent plants per treatment in each experiment.

Next, we assessed the effect of *Pab* infection on leaf cell morphology and organization. Paraffin-embedded cross-sections of infected beet leaves at 15 dpi were differentially stained with safranin and fast green and visualized by light microscopy. We revealed that cross-sections of mock and *Pab*hrcC^−^ treated leaves share similarities, having normal epidermis and mesophyll cells. On the other hand, *Pab*WT-treated leaves demonstrated gall-like structures with ruptured epidermis at random positions. The cross-section of these galls exhibits fast green-stained, non-lignified disorganized small cells, indicating cell hyperplasia (Fig. 3).

Our analyses indicate that *Pab* induces significant tissue deformations by altering mesophyll cell morphology and developmental trajectory, leading to hyperplasia, likely due to uncontrolled cell division.

**Fig. 3.**
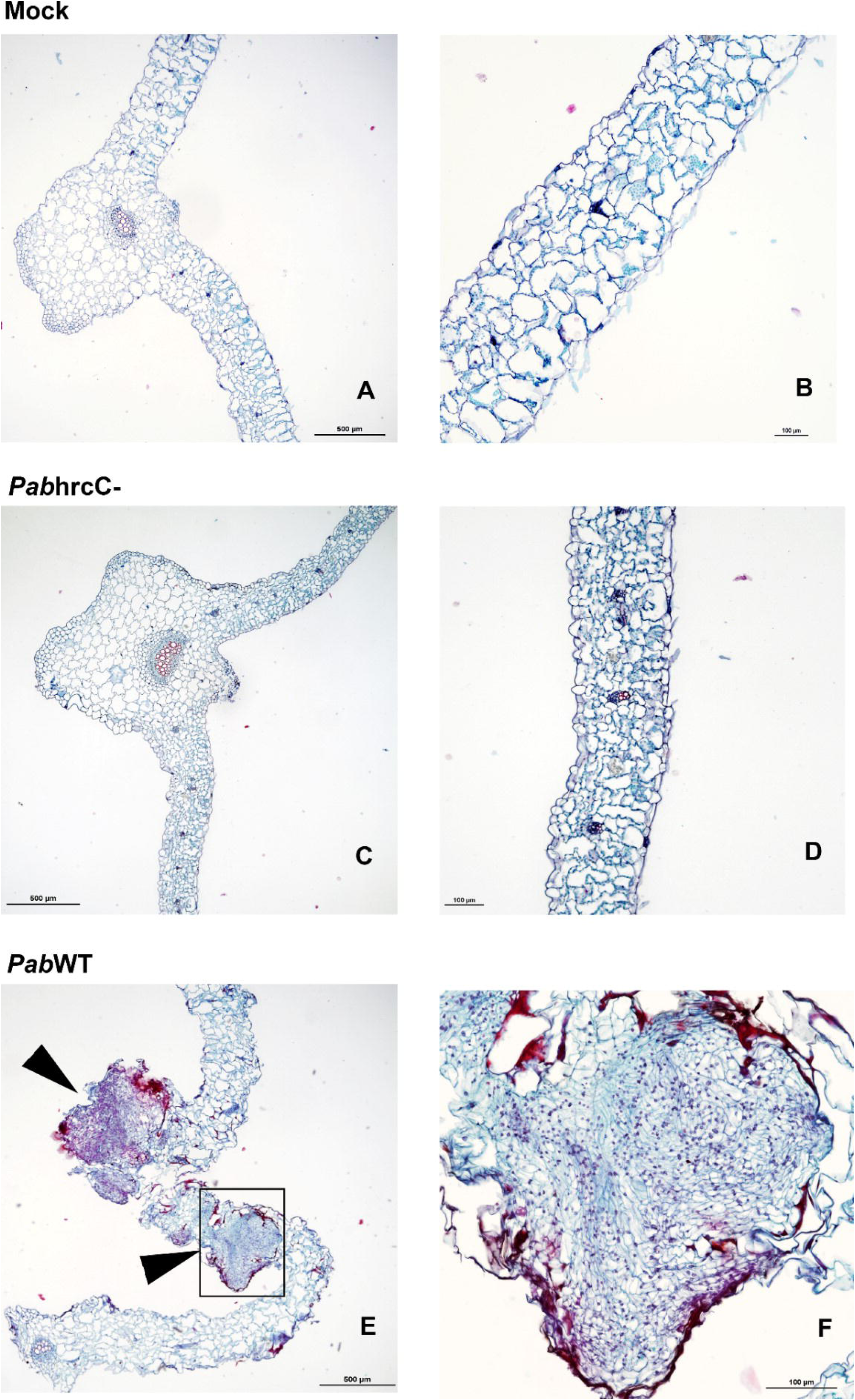
Beet leaf cross section of mock, *Pab*hrcC^−^ and *Pab*WT at 15 dpi. Transverse 10-μm section stained with safranin-fast green for mock (A, B), *Pab*hrcC^−^ (C, D), and *Pab*WT (E, F) were visualized by light microscopy. (E) arrowheads indicate gall formation. (F) Higher magnification of gall tissue showing hyperplasia. Scale bars: 500 μm (A, C and E) and 100μm (B, D and F). The experiments were repeated twice, with five independent plants per treatment in each experiment.

### Gene expression analysis

To uncover the molecular mechanisms underlying the gall formation by *Pab*WT, we compared the expression profiles of mock, *Pab*hrcC^−^, and *Pab*WT-inoculated leaves at 12 and 48 hpi. Three biological replicates were collected after syringe infiltration (10^8^ CFU/ml) of suspensions of *Pab*hrcC^−^, *Pab*WT, or mock, and subjected to RNA sequencing identifying a total of 11,457 genes (FPKM ≥1 cutoff, Supplementary Table 1). Total number of DEGs for different comparisons is shown in Fig. S1A (expression cutoff of FPKM ≥ 1; pFDR < 0.05; fold change difference ≥ 2) (Supplementary Tables 2, 3, 4, 5, 6, and 7). To identify the transcriptomic changes leading to gall formation we compared the transcriptomes of *Pab*WT vs. mock, *Pab*hrcC^−^ vs. mock, and *Pab*WT vs. *Pab*hrcC^−^ infiltrated leaves at each time point. Principal component analysis (PCA) evaluated the replicates of all samples based on the similarity of gene expression profiles (Fig. 4A). Global expression profiles of *Pab*WT and *Pab*hrcC^−^ are more similar to each other after 12 hpi than mock-inoculated samples at 12 hpi. Contrary to that, the *Pab*WT expression profile became very distinct from *Pab*hrcC^−^ and mock at 48 hpi, suggesting specific manipulations done by the T3SS at the transcriptional level (Fig. 4A). In particular, comparative transcriptomic analyses identified 1,755 DEGs (fold change (FC) ≥ 2; pFDR < 0.05) comparing *Pab*WT to *Pab*hrcC^−^ at 48 hpi, including 1,218 up-regulated genes and 537 down regulated genes. Furthermore, venn diagrams were also made based on the up– and down-regulated DEGs (FC ≥ 2; pFDR < 0.05) to show the unique and common genes shared after mock, *Pab*hrcC^−^, and *Pab*WT treatment (Fig 4B-E). For confirmation of RNA sequencing data results, we randomly selected genes from different categories at 12 hpi (6 genes) and 48 hpi (9 genes) and validated their expression using RT-qPCR (Fig. S2 and S3). We found the FPKM values and expression profiles for all the genes after RT-qPCR gave similar results. These results validate that the RNA sequencing data is reliable.

**Fig. 4.**
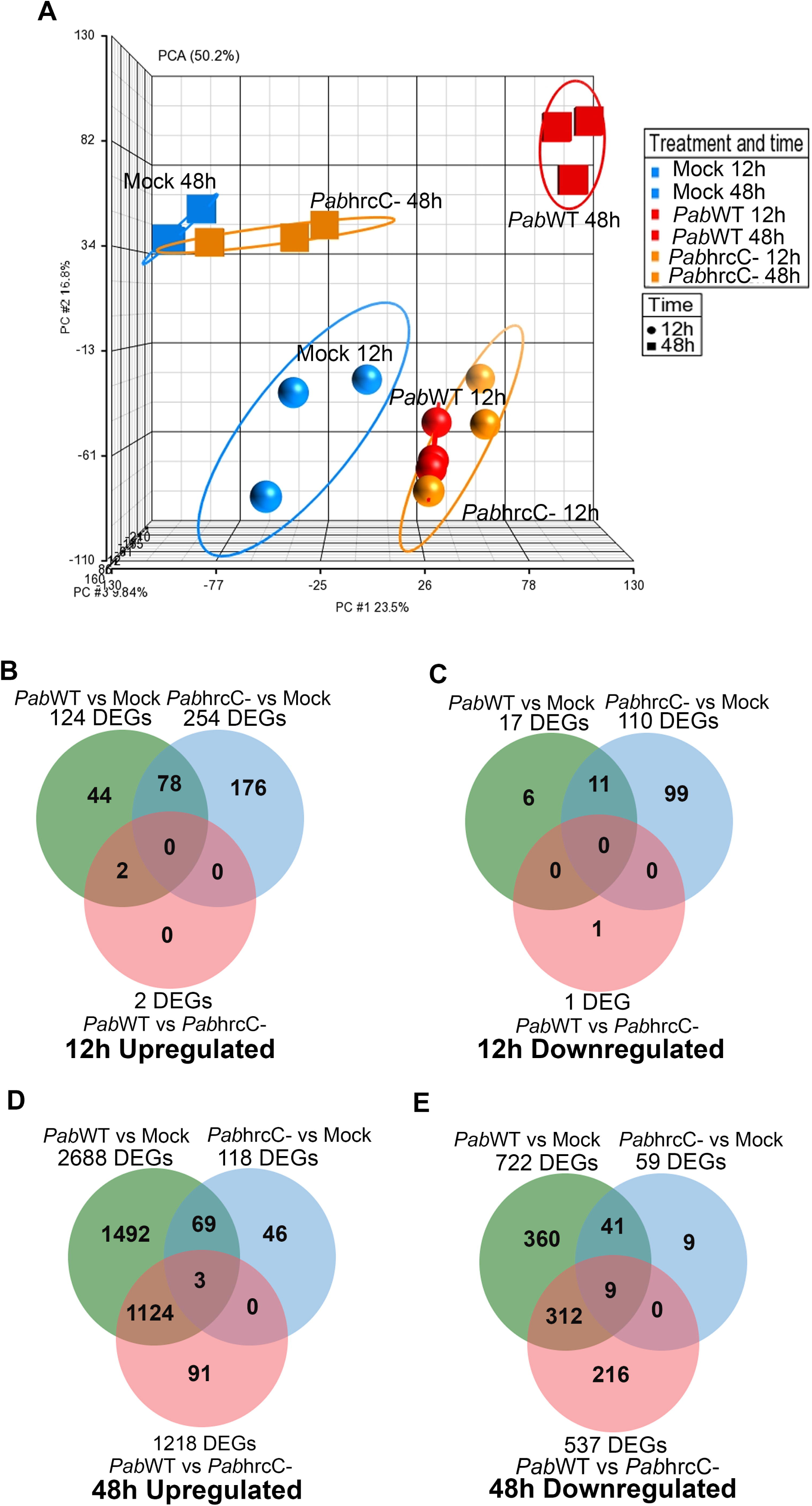
RNA-seq Analysis and Differential Gene Expression in Beet Leaves Post *Pab* Treatments. (A) PCA plots generated using Partek Genomics Suite which represents FPKM values for replicates of RNA-seq samples; blue, red, and orange colour represent FPKM values of different replicates after mock, *Pab*hrcC^−^, and *Pab*WT treatment, respectively. FPKM values ≥ 1 was taken as a cut-off for plotting the data. (B-E) Venn diagram showing the unique and common DEGs among different comparisons, defined as at least a 2-fold change in expression and a false discovery rate (FDR) ≤ 0.05, at 12 hpi (B and C), and 48c:hpi (D and E) in beet leaves.

### *Pab*hrcC^−^ and *Pab*WT induces oxidative stress responsive genes at 12 hpi

To identify the similarities and differences between *Pab*WT vs. mock, *Pab*hrcC^−^ vs. mock, and *Pab*WT vs. *Pab*hrcC-in terms of their transcriptomic profile, corresponding up– and down-regulated gene lists (cutoff of FC ≥ 3) were subjected to GO and KEGG-enriched pathways at 12 hpi and 48 hpi using the STRING database (https://version-11-5.string-db.org) (Szklarczyk et al. 2023). The distribution of these DEGs were allocated to three categories: biological process, molecular function, and cellular compartment (Supplementary Table 8, 9, 10, 11, 12, 13 and 14). Non-redundant top-enriched terms for GO process and function were shown in Fig. 5 (upregulated genes) and supplementary Fig. S1 (downregulated genes).

**Fig. 5.**
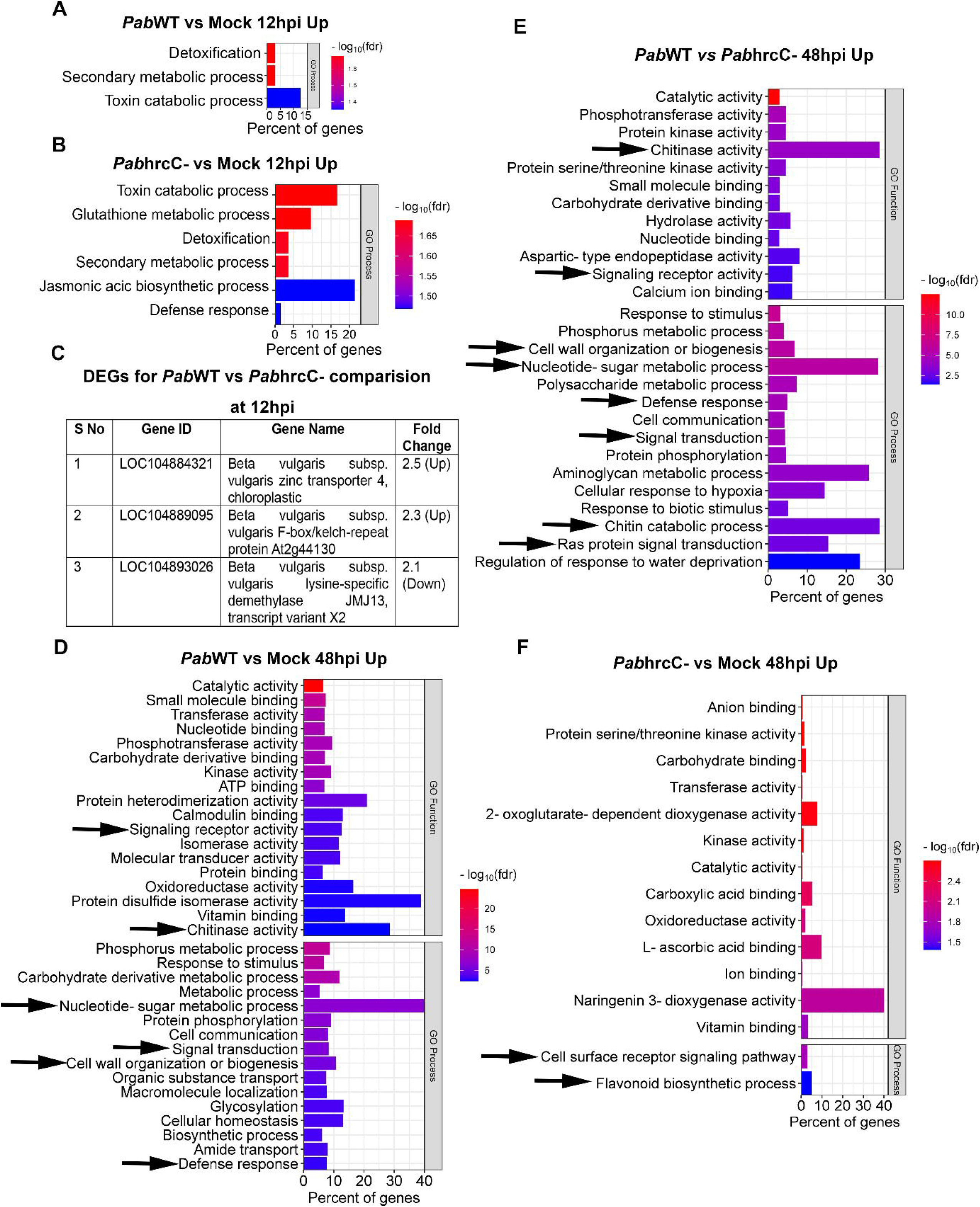
Gene ontology (GO) process and function enrichment analysis of DEGs after mock, *Pab*hrcC^−^, *Pab*WT infiltration in beet leaves at 12 and 48 hpi. GO functional and process enriched categories for upregulated genes: (A) *Pab*WT vs mock at12 hpi, (B) *Pab*hrcC^−^ vs mock treatment at12 hpi, (C) Table for DEGs after *Pab*WT and *Pab*hrcC^−^ at 12hpi, (D) *Pab*WT vs mock at 48 hpi, (E) *Pab*WT and *Pab*hrcC^−^ at 48 hpi. (F) *Pab*hrcC^−^ vs mock treatment at 48 hpi (FDR < 0.05, FC ≥ 3). Black arrows represent important functional groups among different comparisons.

At 12 hpi, we found very few numbers of downregulated genes with a cutoff of FC ≥ 3, which can’t be clustered into GO terms in the *Pab*WT vs. mock and *Pab*hrcC^−^ vs. mock comparisons. Moreover, only three genes were differentially expressed in the case of *Pab*WT vs. *Pab*hrcC^−^ comparison (Fig. 5C; Supplementary Table 4). Among them, the homologs of Arabidopsis genes histone demethylase *JMJ13* was 2.1-fold downregulated, while *zinc transporter 4* and *F-box/kelch-repeat protein At2g44130* were 2.5 and 2.3-fold upregulated, respectively. *JMJ13* repress flowering and also regulates JA and auxin-responsive genes (Zheng et al. 2019; Keyzor et al. 2021). Interestingly, the *F-box/kelch-repeat protein At2g44130* has been known to upregulate in giant cells of root knots and promote nematode susceptibility in Arabidopsis (Curtis et al. 2012).

For upregulated genes at 12 hpi with a cutoff of FC ≥ 3, we found common GO terms related to detoxification, toxin catabolic process, and secondary metabolic process in both *Pab*WT vs. mock and *Pab*hrcC^−^ vs. mock (Fig. 5A and B; Table 1 Supplementary Table 8 and 9). All the three terms include genes encoding glutathione-S-transferases (GSTs) (3 and 4 genes in *Pab*WT vs. mock and *Pab*hrcC^−^ vs. mock respectively) and peroxidases (3 genes in both comparisons), indicating a response against oxidative damage caused by *Pab*WT and *Pab*hrcC^−^ at 12 hpi (Asselbergh et al. 2007; Gullner et al. 2018). Apart from these common terms, we found defense response and JA biosynthetic process terms only in the case of the *Pab*hrcC^−^ vs. mock comparison at 12 hpi (Fig. 5B; Table 1). Defense response term includes genes whose orthologs are known for their antifungal property, SA biosynthetic and signalling genes. *Aspartic proteinase GIP2* (Ma *et al*., 2017) and *endochitinase EP3* (Danaeipour et al. 2023; AJ et al. 1998) are antifungal. Orthologs of SA biosynthetic and signalling genes are *Benzyl alcohol O-Benzoyltransferase* (LOC104902915 and LOC104902913) (Kotera et al. 2023), *PR-10*, and *major allergen Mal d 1* (Puehringer et al. 2003), and *MACPF domain-containing protein CAD1* (Morita-Yamamuro et al. 2005). Altogether, we can conclude that *Pab*hrcC^−^ and *Pab*WT induce genes associated with oxidative damage and defense at 12 hpi, possibly as a result of initial pathogen recognition in the host through PAMP-triggered-immunity (PTI).

**Table 1.**
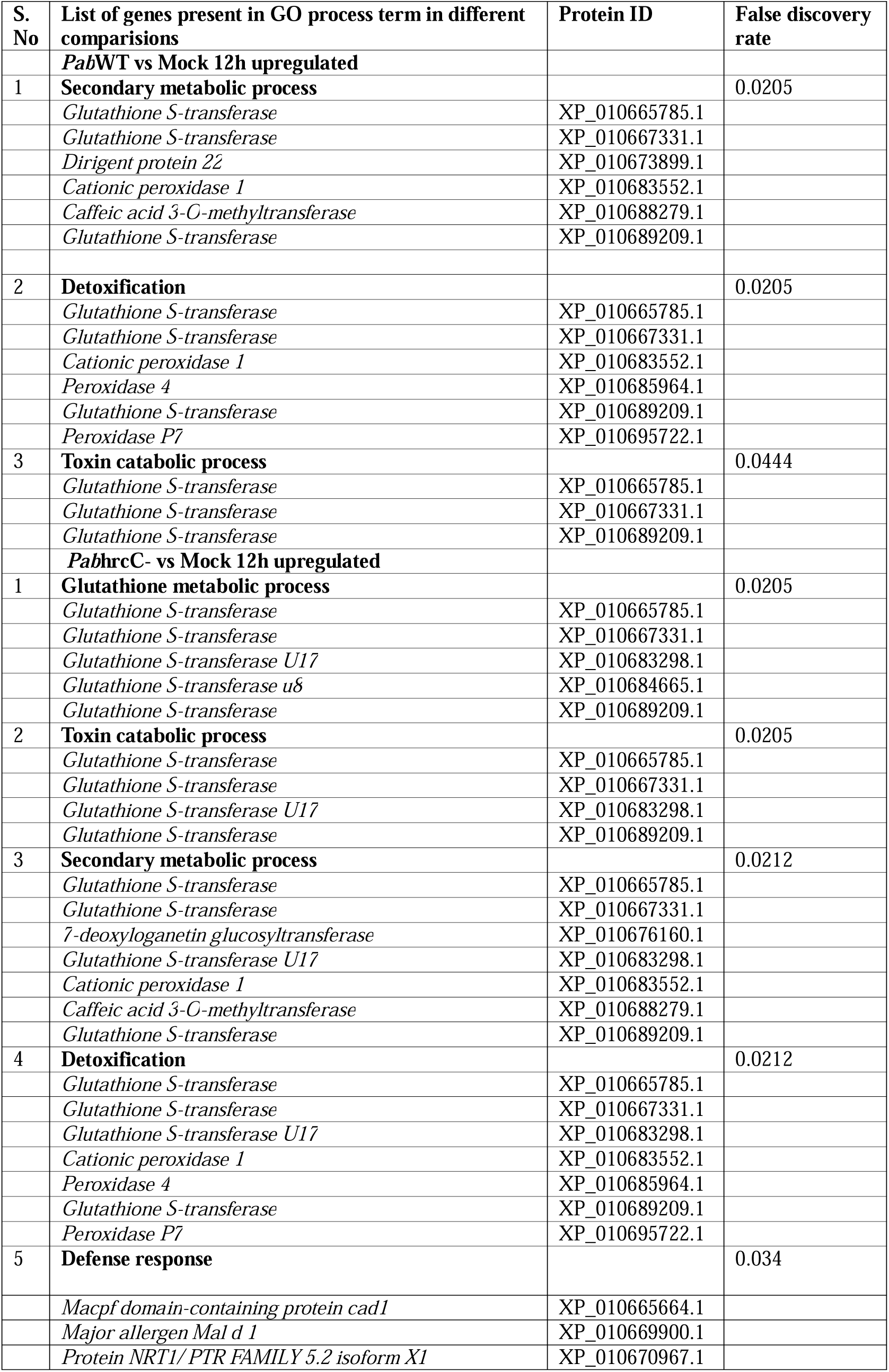
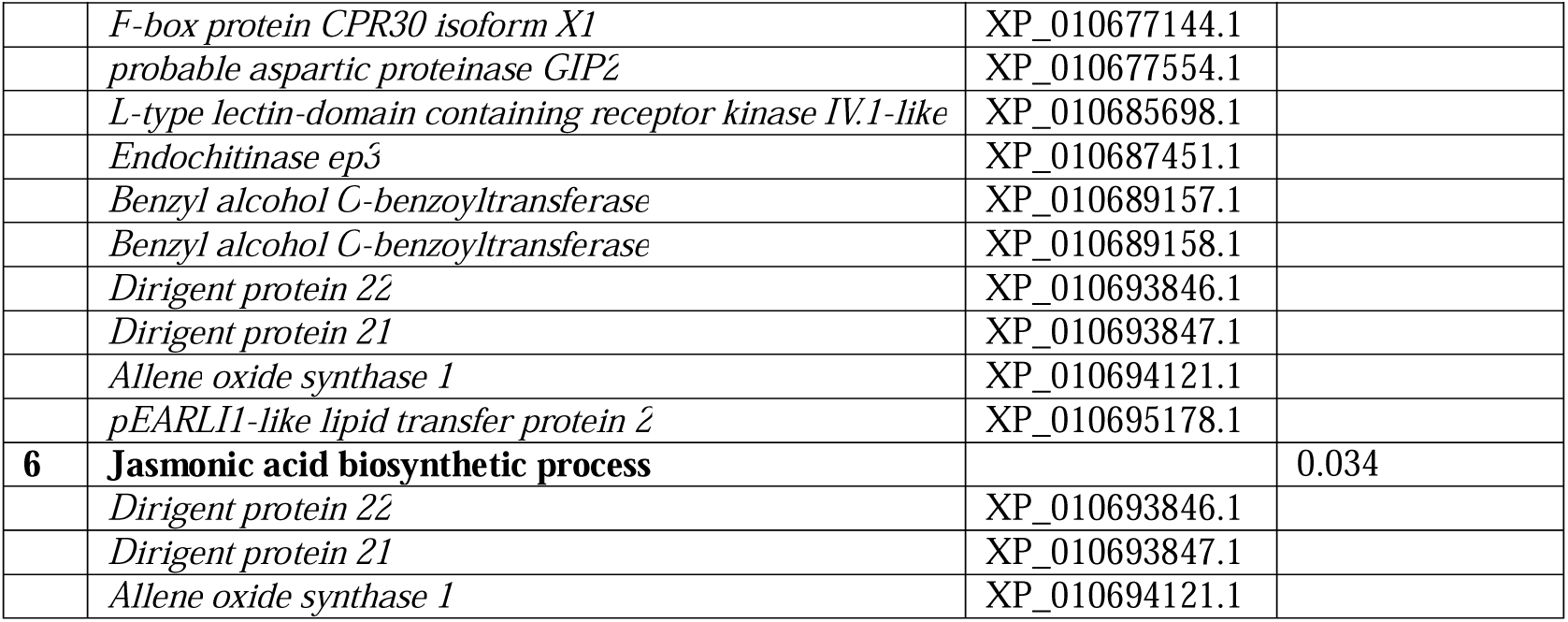
List of upregulated genes found in the functional categories of *Pab*WT vs mock and *Pab*hrcC^−^ vs mock at 12 hpi.

### Cell wall organization related genes are associated with gall formation

To find out the correlation between hyperplasia observed in *Pab*WT-induced galls (Fig 3E and F) and transcriptomic data, we looked into the top GO process terms for upregulated genes (FC ≥ 3) at 48 hpi for all three comparisons, i.e., *Pab*WT vs. mock, *Pab*hrcC^−^ vs. mock, and *Pab*WT vs. *Pab*hrcC^−^ (Fig. 5D, E, and F). *Pab*hrcC^−^ vs. mock comparison revealed only two GO process terms like cell surface receptor signalling pathway and flavonoid biosynthetic process (Fig. 5F). On the other hand, signalling receptor activity, signal transduction, ras protein signal transduction, defense response, nucleotide sugar metabolic process, chitinase activity, and cell wall organization, and biogenesis belong to the top enriched terms and are common in *Pab*WT vs. mock and *Pab*WT vs. *Pab*hrcC^−^ comparisons (Fig 5D and E). Moreover, for downregulated genes (FC ≥ 3) in case of *Pab*WT vs. mock and *Pab*WT vs. *Pab*hrcC^−^ comparison, we found top GO terms like photosynthesis, oxidation reduction process, and chlorophyll biosynthetic process indicating a downregulation of the photosynthetic process due to *Pab*WT inoculation (Fig. S1). We focused our analysis mainly on the *Pab*WT vs. *Pab*hrcC^−^ comparison to find out the genes that are induced by the presence of the T3SS, presumably through the function of T3Es. We hypothesized that genes related to nucleotide sugar metabolic process, chitinase activity, and cell wall organization play essential roles in synthesizing cell wall components or remodelling during cell division and gall formation. We selected genes from these functional categories present in *Pab*WT vs.

*Pab*hrcC^−^ comparison and generated heatmaps, to compare their expression profile among mock, *Pab*hrcC^−^ and *Pab*WT together (Fig. 6). We found expression of all the genes present in these categories were highest in *Pab*WT-treated leaves compared to mock and *Pab*hrcC^−^. This suggests that T3Es, delivered by the T3SSare required to induce genes related to cell wall remodelling or biogenesis.

**Fig. 6.**
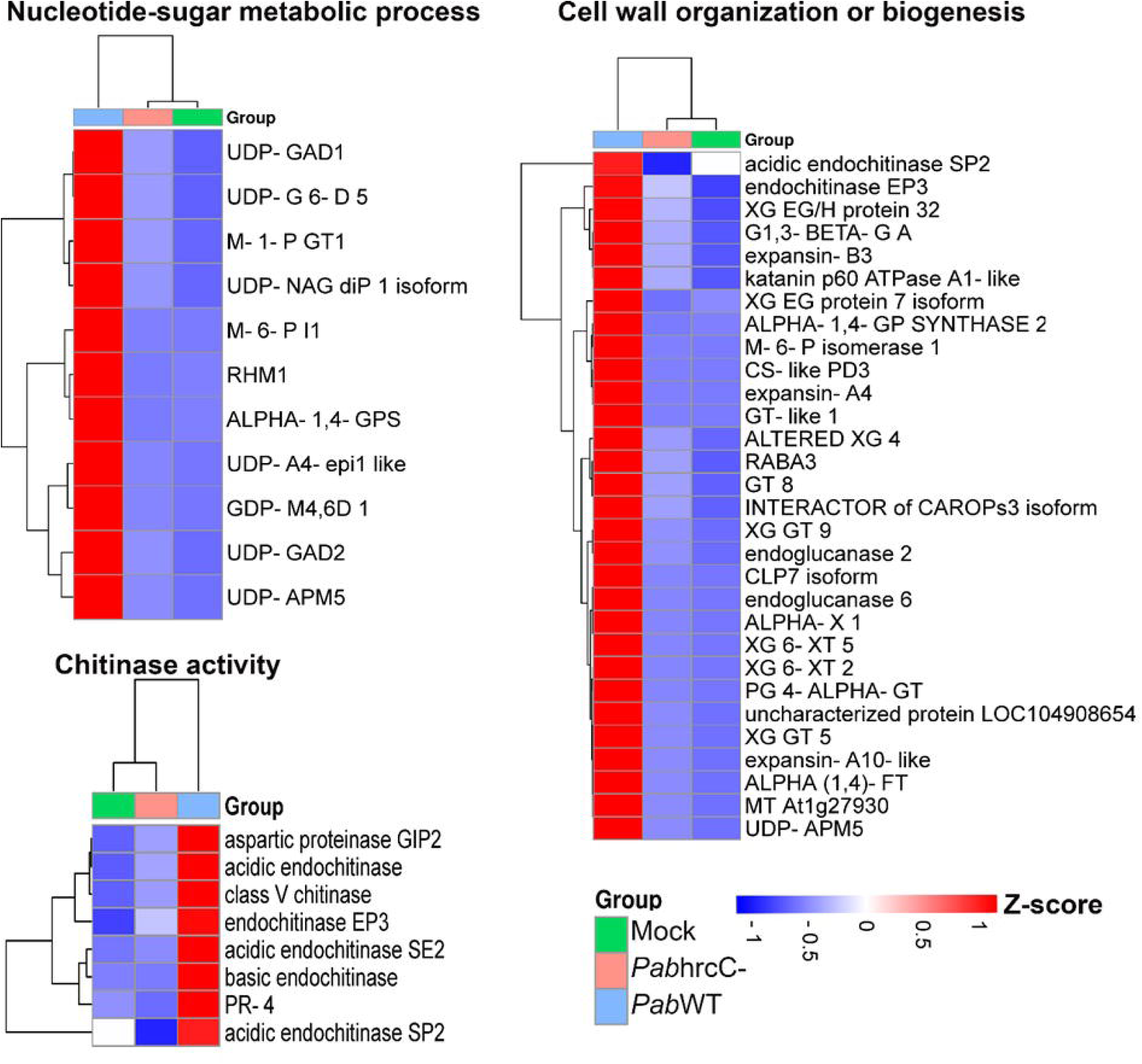
*Pab* induce gene associated cell wall remodelling. Heatmaps for the FPKM (average of replicates) values of the potential upregulated genes (cutoff FC ≥ 3) responsible for cell wall biogenesis or remodelling which belongs to different functional categories for *Pab*WT vs *Pab*hrcC^−^. Details for abbreviation of the gene names are provided in Table 2.

**Table 2.**
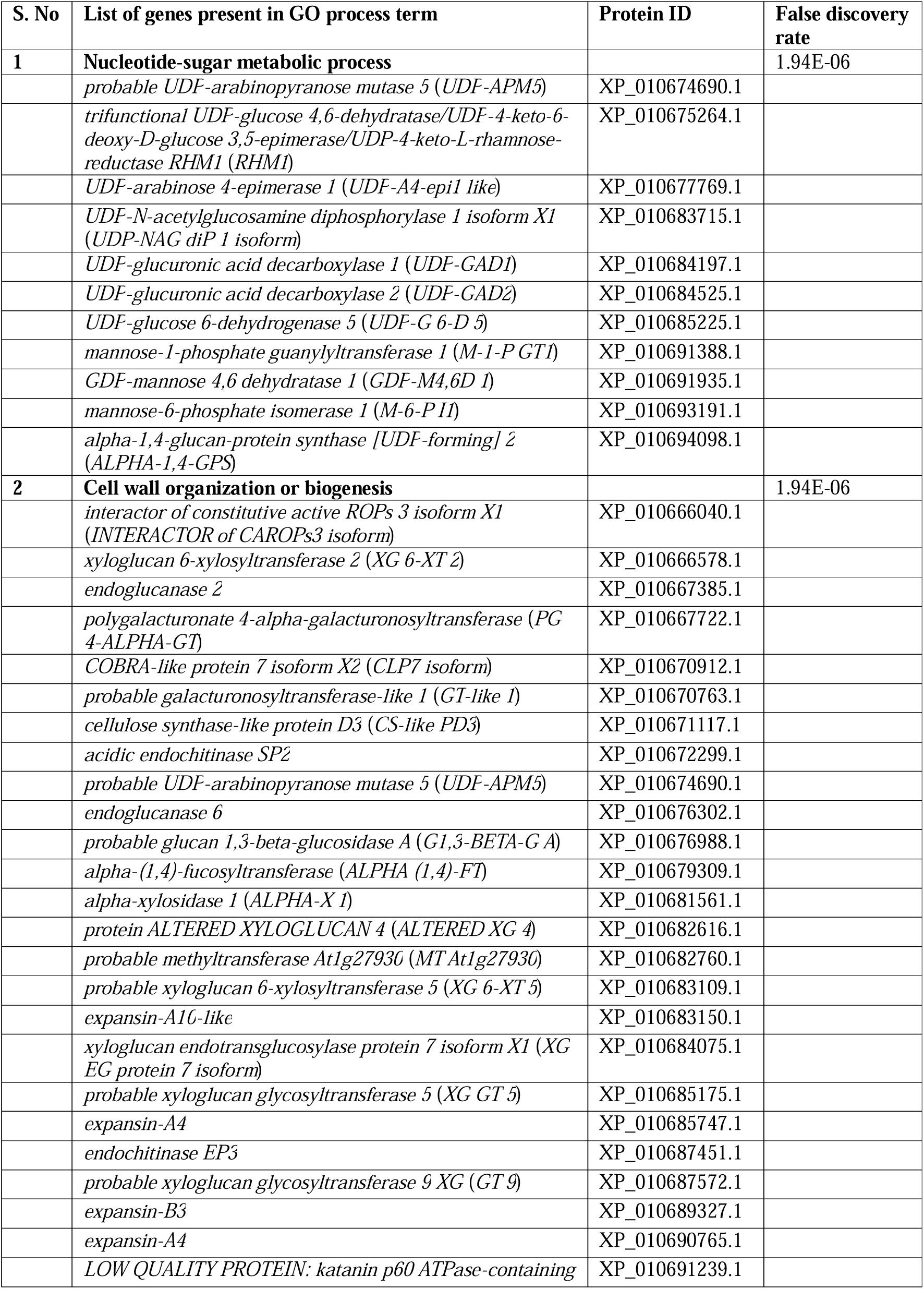
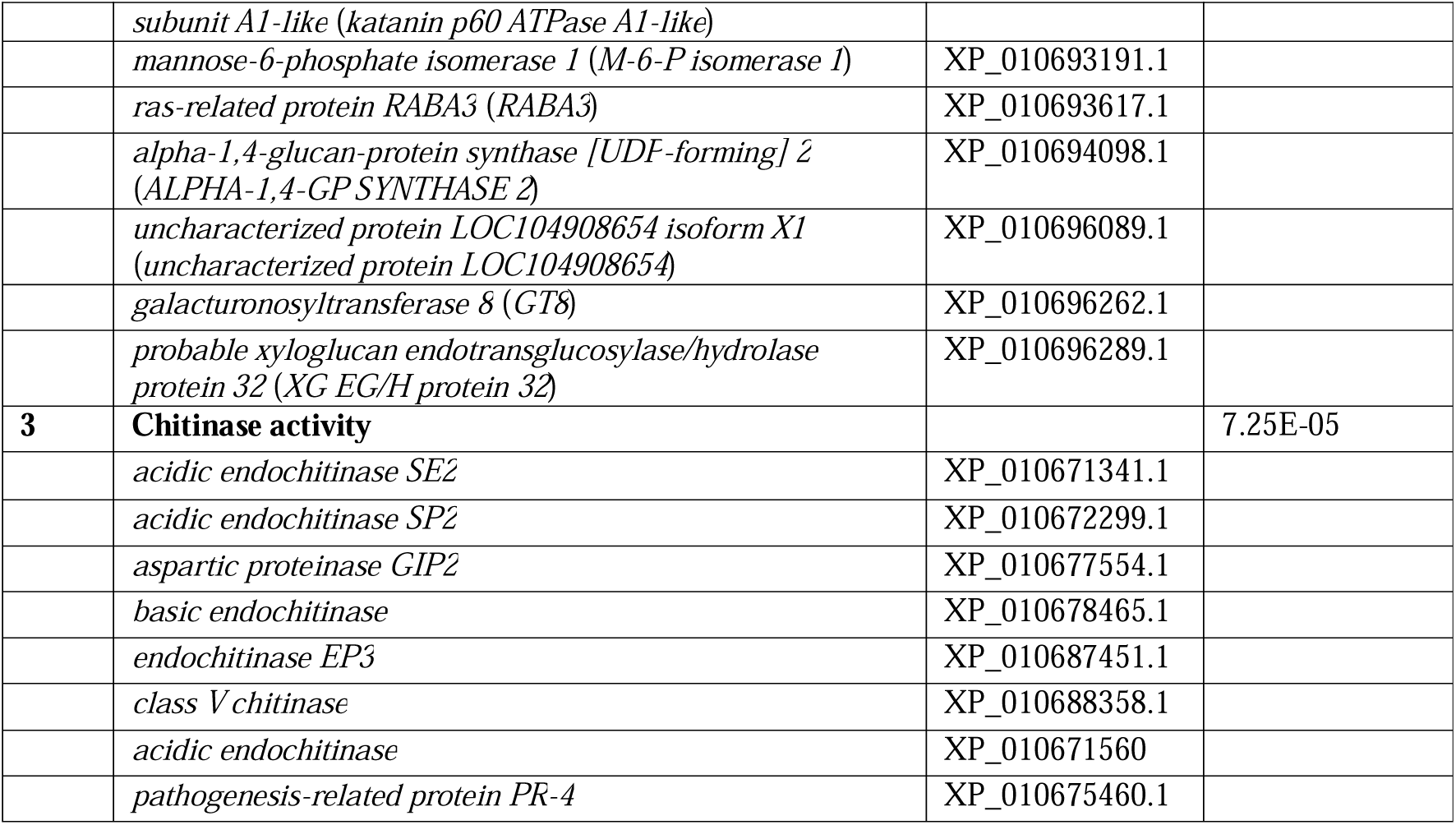
List of upregulated genes found in the functional categories like cell wall organization, nucleotide sugar metabolic process and chitin catabolic process in *Pab*WT vs *Pab*hrcC^−^ comparison at 48 hpi.

We found 31 upregulated genes for cell wall organization in the *Pab*WT vs *Pab*hrcC*-* comparison (Supplementary Table 13; Table 2). Among them some of the interesting homolog genes: *expansin, xyloglucan 6-xylosyltransferase 2*, *cellulose synthase*, *galacturonosyltransferase* are known for cell wall loosening, xyloglucan, cellulose, and pectin biosynthesis respectively (Culbertson et al. 2018; Marowa et al. 2016; Sterling et al. 2006; McNamara et al. 2015). Nucleotide sugars are an important constituent of cell wall components. We found 11 upregulated genes in the *Pab*WT vs *Pab*hrcC^−^ comparision for biosynthesis of nucleotide sugars. These include *UDP-arabinopyranose mutase 5*, *mannose-1-phosphate guanylyltransferase 1* and *UDP-glucuronic acid decarboxylase* etc (Table 2). The homolog of these genes can synthesize L-arabinofuranosyl, GDP-mannose and UDP-xylose respectively that are essential constituents of various cell wall components like hemicellulose, xylan, and xyloglycans (Lukowitz et al. 2001; Konishi et al. 2007; Oka et al. 2007; Kuang et al. 2016; Zhang et al. 2021). Interestingly, we also found 8 genes related to chitinase activity, including *basic endochitinases*, *acidic endochitinase SE2* and *SP2*, *endochitinase EP3*, *class V chitinase*, *PR-4* and *aspartic proteinase GIP2* (Fig 6; Table 2). The homologs of these chitinases are known for their roles in fungal defense and could also play a role in cell division. For instance, *EP3*, *SP2*, *SE2*, *Class V chitinase*, and *aspartic proteinase GIP2* are known for their antifungal property (Danaeipour et al. 2023; Yerzhebayeva et al. 2018; NIELSEN 1993; Ohnuma et al. 2011; Ma et al. 2017b). Also, *basic endochitinases* and *EP3* can help in somatic embryogenesis by modifying arabinogalactan proteins (AGPs) and induce cell division in carrot (AJ et al. 1998; Domon et al. 2000; van Hengel 1998). In our data, we also found upregulated genes encoding for arabinogalactan proteins in the comparison between *Pab*WT and *Pab*hrcC-e.g. LOC104896053 (*arabinogalactan protein 9*) and LOC104891863 (*fasciclin-like arabinogalactan protein 1*) (Supplementary Table 7). It is possible that *Pab*WT might employ similar strategy for inducing cell division during gall formation. Considering the ruptured epidermis observed in gall tissue (Fig 2 and 3), the induction of genes involved in chitinase activity might be crucial for sustaining galls against other microbial infection. Overall, our analysis suggests potential mechanisms by which *Pab*WT manipulates host cellular processes to induce structural changes associated with gall formation.

### *Pab*hrcC^−^ induces betalain accumulation

Symptoms observed in *Pab*hrcC^−^ treated leaves show red coloured betalain accumulation, as shown in Fig. 1A. To further confirm this observation, we quantified betalain components, i.e., betanin and vulgaxanthin-I pigments, spectrophotometrically. Light absorption was measured at 538 nm and 476 nm to calculate the betanin and vulgaxanthin-I concentrations, respectively (von Elbe 2001). We observed the highest amount of both betanin and vulgaxanthin pigment in *Pab*hrcC^−^ treated leaves compared to mock and *Pab*WT treated ones at 10 dpi. Contrary to that, *Pab*WT suppresses vulgaxanthin accumulation compared to mock and *Pab*hrcC^−^ treatments. However, betanin concentration in *Pab*WT-treated leaves was similar to mock treatment (Fig. 7A).

**Fig. 7.**
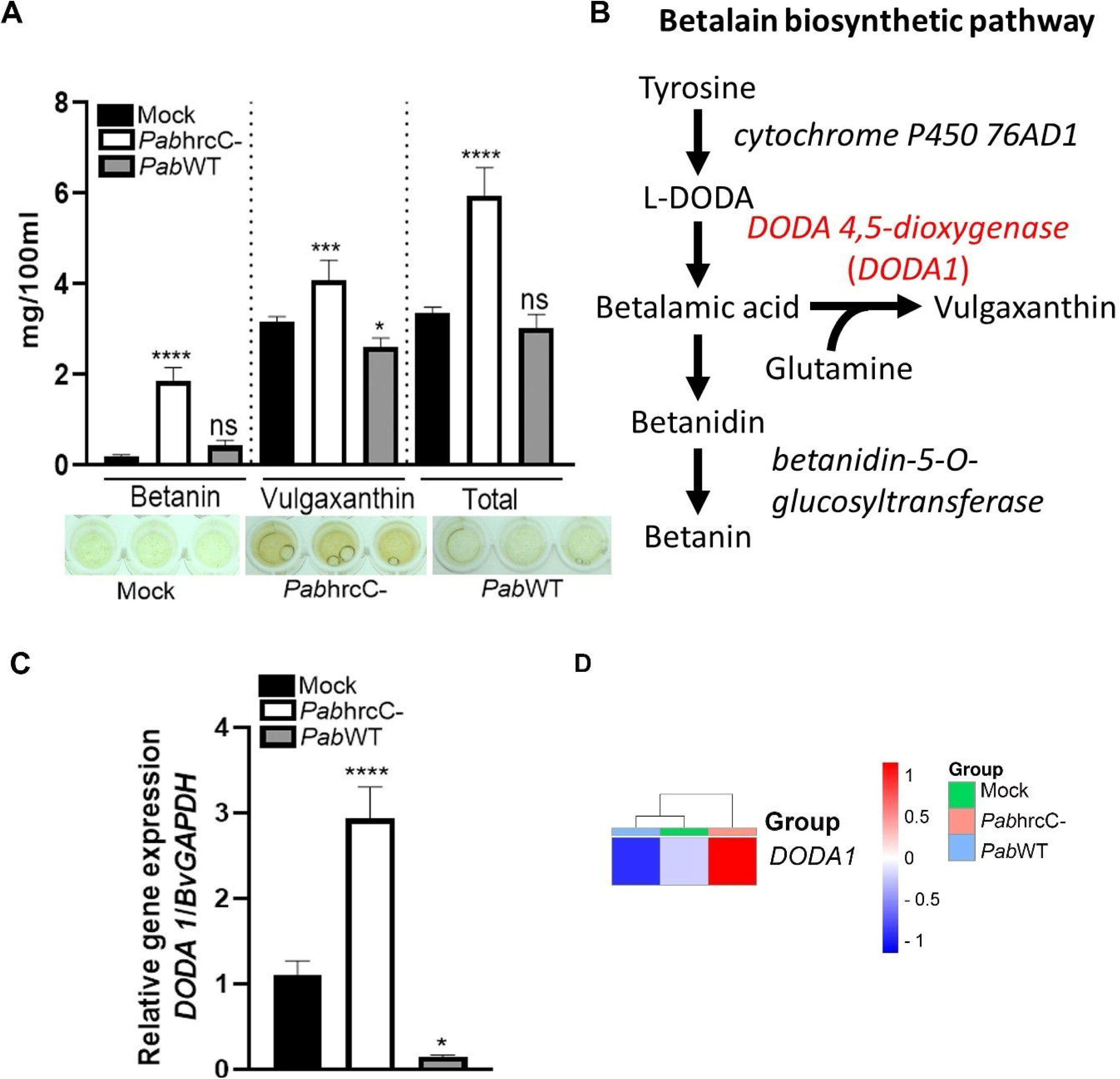
Betalain quantification and relative gene expression of betalain biosynthetic genes. (A) ddH_2_O (mock), *Pab*hrcC^−^ (10^8^ CFU/ml) and *Pab*WT (10^8^ CFU/ml) were infiltration into beet leaves and betalains were quantified at 10 dpi. (B) Schematic diagram to show betalain biosynthetic pathway in beet. (C) Relative gene expression of *DODA1* (*4,5-DOPA DIOXYGENASE EXTRADIOL 1*) after 48 hpi quantified by RT-qPCR. *BvGAPDH* was taken as reference gene. (D) Heatmap of the FPKM (average of replicates) values of betalain biosynthetic gene *DODA1*. The asterisks represents statistically significant difference of the treatments compared to mock based on five (A) or three (C) biological replicates per treatment. It is determined by one-way ANOVA method (Tukey’s multiple comparisons test); * (p≤0.05); ****(p ≤ 0.01). The experiments were repeated twice with similar results.

To examine whether this observation is supported by the transcriptome data, we focused on the expression of the betalain biosynthetic gene *DOPA 4,5-dioxygenase estradiol 1* (*DODA1*) (Polturak and Aharoni 2018; Tanaka et al. 2008) among the three treatments (Fig. 7B). We prepared a heatmap using FPKM values for LOC104908371 (*DODA1*) and performed RT-qPCR at 48 hpi (Fig. 7C and D) and found that *DODA1* gene expression in *Pab*hrcC^−^ inoculated leaves was induced compared to mock and *Pab*WT treatments. Contrary to that, *Pab*WT suppresses *DODA1* expression compared to mock. These results supports the betanin quantification results as found in Fig. 7A. A similar gene expression profile for *DODA1* using RT-qPCR and FPKM values validates our RNA sequencing data results (Fig. 7C and D).

These results suggest that betalain accumulates in beet leaves as a response to *Pab* colonization. However, the presence of a functional T3SS suppresses this response, likely through the action of T3SS effectors, which reduce the transcript accumulation of *DODA1*.

## Discussion

In this study, we characterized the contribution of *Pab* T3SS to gall formation in beet leaves and identified key regulatory pathways induced by *Pab* at early infection on the onset of gall formation. We identified that *Pab* induce gall-like structures on root and leaves, which are associated with cell hyperplasia and tissue ruptures, in dependence on the T3SS. In addition, comparative transcriptome analyses identified major transcriptional reprograming mediated by *Pab* T3SS effectors as of 48 h post inoculation, long before gall formation is visually apparent, which includes the induction of multiple pathways associated with signal transduction, defense, cell wall modulation, and suppression of betalain pigment production.

To achieve uniformed tissue exposure to *Pab*, we standardized leaf inoculation assays by syringe infiltration. Our analyses confirmed that *Pab* can promote gall formation on leaf tissue as well, and, similarly to root inoculation, gall formation is eliminated when leave are inoculated with a mutant in the T3SS apparatus, indicating that its dependant on delivery of T3SS effectors (Geraffi et al. 2023). Additionally, wild-type bacteria demonstrate 7 to 8-fold increase in leaf bacterial population and significant reduction in betalain accumulation compared to the T3SS mutant strain.

The contribution of T3SS effectors to bacterial colonization is well documented in numerous plant pathogenic bacteria and is typically associated with suppression of plant immune responses, alteration of plant metabolite homeostasis, cell identity, or water transport (Schreiber et al. 2021). *Pab* encodes for several T3SS effectors that share high homology to *Pseudomonas syringae* and *Erwinia amylovora* T3SS effectors associated with manipulation of immunity signaling and increase in apoplast water availability such as HopX1, HopAF1, and DspE (Nomura et al. 2024; Gimenez-Ibanez et al. 2014; Castañeda-Ojeda et al. 2017; Geraffi et al. 2023). Therefore, the increase in *Pab* leaf colonisation can be attributed to other pathways associated by T3SS effectors in addition to the environment produced within the gall. Interestingly, despite the clear reduction in leaf population, the *Pab* T3SS mutant can still reach relatively high bacterial populations compared to T3SS mutants of other pathogens such as *Xanthomonas*, *Ralstonia* and *Pseudomonas syringae* sp. (Schreiber et al. 2021), implying that *Pab* does not significantly depend on T3SS effectors for host colonization. This data aligns which previous reports of non-pathogenic *P. agglomerans* as a dominant plant endophyte that demonstrates high variance in the presence or absence of a T3SS, and the phyletic lineages of the T3SS, when present (Moretti et al. 2021; Walterson and Stavrinides 2015), and supports the hypothesis that *Pab* is a recently adapted pathogen developed from commensal *P. agglomerans* (Barash and Manulis-Sasson 2009).

Another notable phenotype observed in the comparison between wild-type *Pab* and T3SS mutant *Pab* is the suppression of leaf betalains accumulation in the infiltrated areas. Betalains are nitrogen-containing pigments uniquely found in *Caryophyllales*, which have been attributed with traits such as scavenging of oxidative radicals and antibacterial activity (Gliszczyńska-Świgło et al. 2006; von Elbe 2001). Accordingly, they are hypothesized to protect *Caryophyllales* from abiotic and biotic stress. Betalain accumulation by the *Pab* T3SS mutant but not *Pab*WT, suggests that these pigments might play an intrinsic role in host basal defense against pathogens, which is supressed by T3SS effectors (Schreiber et al. 2021). This pigment accumulation can be a direct response to pathogen attack or a secondary response to oxidative stress activated during pathogen recognition. Considering that suppression of betalains accumulation is accompanied by down regulation of *DODA1*, which play a key role in betalain biosynthesis (Schwinn 2016), the T3SS suppression of this pathway is likely to be associated with manipulation of pathogen sensing or signaling, rather than active biochemical manipulation of betalain biosynthesis or degradation. In addition, the JA biosynthesis pathway, typically linked to defense, is induced by the *Pab* T3SS mutant but not by *Pab*WT at 12 dpi. It would be valuable to investigate the mechanism by which the T3SS influences betalain regulation, whether this response is JA-mediated, and which T3SS effectors target this pathway, along with the underlying targeting mechanisms, in future studies. *Pag* induced galls have been previously subjected to histological and biochemical characterization in gypsophila stem cuttings (Chalupowicz et al. 2006). These galls, which were in large amorphous sponge-like structures, were composed of large expended cells surrounded by clusters of small undifferentiated cells undergoing cell division (Chalupowicz et al. 2006), indicating that *Pag* induce both hypertrophy and hyperplasia. This cellular composition is somewhat dissimilar of what we observed in beet leaf tissues, in which gall structures were solely composed by small undifferentiated cells. These discrepancies can be a result of differences in host, target tissue, and pathogen. Previous findings have shown that *Pag* and *Pab* galls in gypsophila stem cuttings are dissimilar in structure and pigmentation (Barash and Manulis-Sasson 2007). These differences might be a result of differential arsenal of T3SS effectors associated with gall development between the two pathogens (Manulis and Barash 2003; Geraffi et al. 2023), which may target different cellular pathways. In addition to pathogen effector arsenals and host, the differences in cell differentiation within the galls could be a result of target tissue. Indeed, gall morphology and the developmental kinetics differ greatly between beetroot and beet leaves, as root galls are more robust and defined, and appear earlier compared to leaf galls that appear later, smaller and more localized. This suggests that, outside of direct reprogramming by pathogen effectors, tissue structure, biochemistry and physiology play a role in the gall developmental trajectory.

To decipher the mechanisms that facilitate tumor development by *Pab*, we conducted comparative transcriptome analyses between beet leaves infected with wild type *Pab* to *Pab hrcC* mutant that cannot promote gall formation. We set to identify genes associated with gall initiation and, therefore, conducted RNA sequencing at the early stages of infection, before gall is apparent, at 12 and 48 hpi. Transcriptome analyses revealed that significant transcriptional reprograming by *Pab* T3SS effectors are apparent as of 48 hpi associated with close to 2,000 DEGs. GO enrichment analyses identified multiple pathways associated with the onset of gall formation including up-regulation of cell wall remodeling, carbon metabolism, and signal transduction and down regulation of photosynthesis, suggesting major shifts in cell structure and energy homeostasis. Multiple genes associated cell wall loosening are significantly elevated by *Pab*, these include expansins, endoglucanases, and polygalacturonase (Cosgrove 2016). Cell wall loosening is important for remodeling cell architecture, and with coordination of cell cycle control, is essential for cell division (Sablowski and Carnier Dornelas 2014). Indeed, up-regulation of cell wall loosening associated genes was reported in transcriptome analyses of pathogen-induced hyperplasia by *Agrobacterium tumefaciens* during crown gall development or by *Xanthomonas citri* during pustule formation (Tkachenko et al. 2021; Hu et al. 2016; Zou et al. 2021; Deeken et al. 2006). Expansins, in particular, have been reported to be essential for gall and pustule formation during crown gall and citrus canker disease (Anand et al. 2007; de Souza-Neto et al. 2023). However, hyperplasia associated transcriptome of both pathogens also showed significant increase in gene associated with DNA replication, cell cycle control and cell proliferation (Deeken et al. 2006; Zou et al. 2021; Hu et al. 2016), which were not enriched in *Pab*-inoculated beet leaves. This result is surprising, considering that *Pab* leaf galls are associated with unstructured small cells, suggesting uncontrolled cell division and not cell expansion, which is expected in the case that cell wall loosening associated genes are up regulated and cell division and DNA replication associated genes are not. We note that *Pab*-mediated galls are only apparent as of 10-15 days post inoculation, which is 8-13 days after the time point used for transcriptome analysis. It is possible that cell division associated pathways are induced at a later stage of gall development, as a secondary response to changes in cell wall architecture combined with other signals. We note that cell wall rearrangement also occurs during the activation of plant defense responses (Wan et al. 2021). It is possible that an increase in cell wall-associated enzymes might also be linked to alterations in defense responses induced by an increase in PAMP elicitors due to the higher bacterial titre of the *Pab*WT or the induction/suppression of immune responses associated with T3Es (Schreiber et al. 2021). However, considering that we did not observe an increase in cell wall-associated genes in the *Pab* T3SS mutant, compared to mock treatment, this possibility is less likely since the PTI response is also induced in this scenario. Up-regulation of numerous genes associated with carbohydrate metabolism, and the down regulation of photosynthesis genes by *Pab*, compared to the *hrcC* mutant, suggest potential manipulation of sink source dynamics, altering beet leaves into a source tissue, and thus, securing, nutrient for the pathogen within the gall (Harris and Pitzschke 2020). However, it should be noted that these manipulations are not unique to gall-associated pathogens and can be found in many pathosystems, and are transitionally considered to be more associated with growth-defense trade-offs than altered sink source dynamics (Huot et al. 2014). In addition, altered carbohydrate profile might serve as a signal for cell differentiation. Indeed, carbohydrate metabolism have long been associated with differentiation and cell identify, either by affecting metabolic homeostasis, regulating phytohormones or by directly acting as signal molecules or their precursors (Eveland and Jackson 2012; De Coninck et al. 2021).

While our study provided initial indications of cellular physiological changes associated with the early onset of *Pab*-mediated gall formation, the molecular mechanism that initiates it is still elusive. Previous studies conducted on *Pag* in gypsophila reported that the manipulation of cell identity to produce gall is facilitated by the activity of two effectors, PthG and HsvG, that may work in cooperation with one another, potentially through modulation of auxin accumulation and/or polar transport by an unknown mechanism (Weinthal et al. 2010; Nissan et al. 2019; Chalupowicz et al. 2013). While gene groups associated with auxin metabolism, sensing, and transport were not enriched in the beet transcriptome, cell wall loosening genes, which are typically associated with auxin response (Perrot-Rechenmann 2010), are induced. This suggests that *Pab* effectors might manipulate auxin response as well, but this manipulation may be either post-translational, too fine-tuned to be detected in the transcriptome, or act downstream of auxin sensing. Considering that gall formation is mediated by a small number of effector proteins, it is expected that these effectors will target key host factors, initiating a chain of events leading to gall formation in a manner similar to the direct transcriptional induction of the *CsLOB1* susceptibility gene by the transcriptional activator-like effector (TALE) PthA4 of *Xanthomonas citri* (Hu et al. 2014). Similarly to PthA4, HsvG and HsvB have DNA binding and eukaryote transcriptional activation activity (Nissan et al. 2006), suggesting that they function as transcriptional regulators in the host. Considering that HsvG and HsvB significantly contribute to gall formation on gypsophila and beet (Nissan et al. 2006; Valinsky et al. 2002), it is likely that these effectors promote the accumulation of susceptibility targets as well. However, unlike PthA4, which belongs to the well-studied TALE family (Teper et al. 2022), the DNA codon affinity and transcriptional activation mechanisms of HsvG and HsvB are still elusive, and their targets cannot be predicted through computational analysis of host promoter regions. The comparative beet transcriptome conducted in this study may be used as a tool to identify HsvB susceptibility targets in future functional assays.

In conclusion, this study characterized the cellular architecture and transcriptional profile of *Pab*-mediated galls on beet leaves. The results indicate that *Pab*-mediated gall formation in beet leaves depends on the delivery of effector proteins by the T3SS and is facilitated by large-scale transcriptional reprogramming, resulting in tissue hyperplasia through alterations in cell wall architecture and manipulation of sink-source carbon metabolism. These findings provide crucial insights into the gene pathways that facilitate cellular reprogramming in bacterial galls targeted by secreted bacterial virulence factors.

## Material and Methods

### Plant growth conditions, bacterial strains and pathogenicity test

WT and type III secretion system mutant (hrcC^−^) of *Pab* strain 4188 has been used in this study (Geraffi et al. 2023; Burr et al. 1991; Geraffi 2021). Bacteria were grown in Luria Bertani (LB) broth or LB agar plates with 100 µg/ml rifampicin at 28°C.

Seeds of beet (*Beta vulgaris*) cv. Red Egyptian were germinated for 10 days. The seedlings were then transferred into pots and grown in a growth chamber at 21 °C with a photoperiod of 12 h. Mature leaves from 1.5-month old beet plants were infiltrated with mock (ddH_2_O), suspension (10^8^ CFU/mL) of *Pab*hrcC^−^, or *Pab*WT. Symptoms were photographed on 10 dpi and 20 dpi. To determine leaf bacterial growth, five samples of four leaf discs (1 cm diameter) were collected from five plants at 0, 5, 10, and 15 dpi and homogenized in 1 ml of ddH_2_O and CFU were counted via serial dilution plating.

Pathogenicity tests on table beet cubes based on (Ezra et al. 2000, 2004) were performed with some modification. Briefly, mature beetroots were soaked in 1% sodium hypochloride for 10 min and then washed twice in sterile water. They were then cut into small cubes, each of which was placed on sterile 1.5% water agar in a petri dish. Overnight grown bacterial culture on LB Broth was diluted to 10^8^ CFU/mL. Inoculation was done by streaking the bacteria with the help of sterile toothpick dipped in 10^8^ CFU/mL bacterial culture. The petri dishes were incubated for 6 days at 21°C in the growth room and then tested and photographed for gall formation. Three beet cubes were used for each strain, and the experiment was performed at least three times.

### Scanning electron microscopy (SEM), paraffin embedded sectioning of leaves and microscopic visualization

For SEM, samples from syringe infiltrated sites from beet leaves were collected after 10 and 20 dpi. These samples were fixed in 2.5% glutaraldehyde in PBS. After washing with PBS, samples were dehydrated by successive ethanol dehydration (25%, 50%, 70%, 90% and 100% EtOH each for 30 min). After critical point drying (Balzer’s critical point drier) the samples were mounted on aluminium stubs and sputter-coated (SC7620, Quorum) with gold. Images were captured on the scanning electron microscope (JCM-6000, JEOL). For microscopic visualization of leaf and gall sections, the samples were collected after 15 dpi. Samples were fixed for 24 h in formaldehyde: acetic acid: ethanol 70%, 10:5:85, v/v (FAA) solution. Fixation was followed by an ethanol dilution series (50, 70, 95,100%; 1 h each and 100% overnight) and a subsequent stepwise exchange of ethanol with histoclear solvent (Xylene substitute). Samples were embedded in paraffin and cut by rotary microtome (Leica RM2245) into 10 μm sections, then deparaffinized by histoclear solventfor 12 min 2 times, and rehydrated (ethanol 100, 95, 70, and 50%, 2 min each). Then, samples were stained with Safranin O (1% w/v) for 1 hour, washed with tap water, dehydrated in ethanol (50, 70, and 95%; 30 s each), and stained with fast green (0.3% in 95% EtOH 1:1 clove oil) for 1 min. The stained samples were examined with a light microscope (Nikon eclipse NiE) at 4X and 10X magnifications. Photos acquired by Nikon DS-fi1 camera and NIS elements software.

### Betalain quantification

Betalain was quantified spectrophotometrically (Biotek Synergy HT Multi-Mode Microplate Reader). Briefly, 200 mg of beet leaves was crushed in 500µl of ddH_2_, centrifuged at 13000 rpm for 5 min at 4°C and collected the supernatant. Using 0.05 M phosphate buffer, pH 6.5 as the solvent blank, spectrophotometer was set zero at 476, 538, and 600 nm. Beet leaf extract was diluted with 0.05 M phosphate buffer, pH 6.5 such that the A538 of the sample OD reaches between 0.4 and 0.5. Recorded the absorbance for all samples at 476, 538, and 600 nm. Betanin and vulgaxanthin quantity was calculated as described in (von Elbe 2001)

### RNA isolation, cDNA library construction, RNA sequencing and data analysis

Total RNA was isolated with the RNeasy Plant Mini Kit (QIAGEN) from mock, *Pab*hrcC^−^, and *Pab*WT inoculated leaves at 12 hpi and 48 hpi (three biological replicates for each treatment, in total 18 samples were collected). Total RNA was given to BGI genomics company at China for library preparations and RNA sequencing analysis. Briefly, RNA integrity was evaluated using Agilent 2100 Bioanalyzer. Oligo dT beads were used to enrich mRNA with poly A tail. RNA fragmentation and first strand cDNAs were generated using random N6-primed reverse transcription, followed by the second strand cDNA synthesis. The synthesized cDNA was subjected to end-repair and then was 3’ adenylated. Adaptors were ligated to the ends of these 3’ adenylated cDNA. PCR amplification was performed to enrich the purified cDNA template using PCR primer. PCR product was denatured and cyclized by splint oligo and DNA ligase. DNA nanoball was synthesized and sequencing on DNBSEQ (DNBSEQ Technology) platform was done.

rRNA reads, low quality reads, adaptors and reads with unknown bases (N>5%) were filtered using SOAPnuke software (version: v1.5.2, parameters: –n 0.001 –l 20 –q 0.4 –A0.25, website: https://github.com/BGI-flexlab/SOAPnuke). After reads filtering, mapping was performed to reference genome using HISAT2 (Version: v2.0.4 Parameters: –-phred33 –-sensitive –-no-discordant –-no-mixed –I 1 –X1000 –-rna-strandness RF Website: http://www.ccb.jhu.edu/software/hisat) (Kim et al. 2015). On average 85.62% of the reads are mapped. StringTie (Version: v1.0.4, Parameters: –f 0.3 –j 3 –c 5 –g 100 –s 10000 –p 8 –rf, Website: http://ccb.jhu.edu/software/stringtie) (Pertea et al. 2015) was used to reconstruct transcripts, and with genome annotation information we identify novel transcripts by using Cuffcompare (Version: v2.2.1, Parameters: –p 12, Website: http://cole-trapnell-lab.github.io/cufflinks) (Trapnell et al. 2012) and predict the coding ability of those new transcripts using CPC (Version: v0.9-r2, Parameters: Default, Website: http://cpc.cbi.pku.edu.cn) (Kong et al. 2007) In total, we identify 15,466 novel transcripts. Novel coding transcripts were merged with reference transcripts to get complete reference, then we mapped clean reads to it using Bowtie2 (Version: v2.2.5, Parameters: –q –-phred33 –-sensitive –-dpad 0 –-gbar 99999999 –-mp 1,1 –-np 1 –-score-min L,0,-0.1 –I 1 –X1000 –no-mixed –-no-discordant –p 1 –k 200, Website: http://bowtie-bio.sourceforge.net/Bowtie2/index.shtm) (Langmead and Salzberg 2012). Normalized gene expression level (FPKM) was calculated for each sample with RSEM (Version: v1.2.12, Parameters: default, Website: http://deweylab.biostat.wisc.edu/ RSEM) (Li and Dewey 2011). For 18 cDNA samples, a total of 172.76 GB of clean reads was generated (Supplementary Table 15). Differential gene expressions were detected with DEseq2 (Parameters: Fold Change >= 2.00 and Adjusted P value <= 0.05) (Love et al. 2014). PCA plot (Fig 4) and heatmaps (Fig S1) were generated using Partek Genomics Suite (https://www.partek.com/partek-genomics-suite/; 7.20.0831). Venn diagrams were generated using Venn diagram genomics tool (https://www.biotools.fr/misc/venny). Gene Ontology (GO) and KEGG pathway analysis was performed using STRING tool (https://version-11-5.string-db.org/). SR plot was used to generate GO plots and heatmaps shown in Fig 5 and Fig 6 respectively (Tang et al. 2023). Transcriptomics information of the experiment are available in GEO database (accession number GSE269454).

### RT-qPCR analysis of gene expression

A High-Capacity cDNA Reverse Transcription Kit (Applied Biosystems) (Cat No. 4368814) was used to reverse transcribe 1 µg of RNA samples. Fast SYBR Green Master Mix (Applied Biosystems) was used to amplify cDNAs using gene-specific primers (Supplemental Table

16). The reaction was performed in QuantStudio 1 Real-Time PCR System (Applied Biosystem). The *BvGAPDH* gene was used as a reference gene for normalization, and gene expression was calculated by the comparative Ct method (Pfaffl 2001).

## Acknowledgments

This research was supported by the Israel Science Foundation (ISF) under grant number 488/19. This manuscript is dedicated to the memory of Guido Sessa, a beloved colleague, professor, and friend.

