## Supplementary material for "Comparative transcriptomic profile reveals candidate genes manipulated by type III effectors of *Pantoea agglomerans* pv. *betae* leading to gall formation in beet": Fig. S1

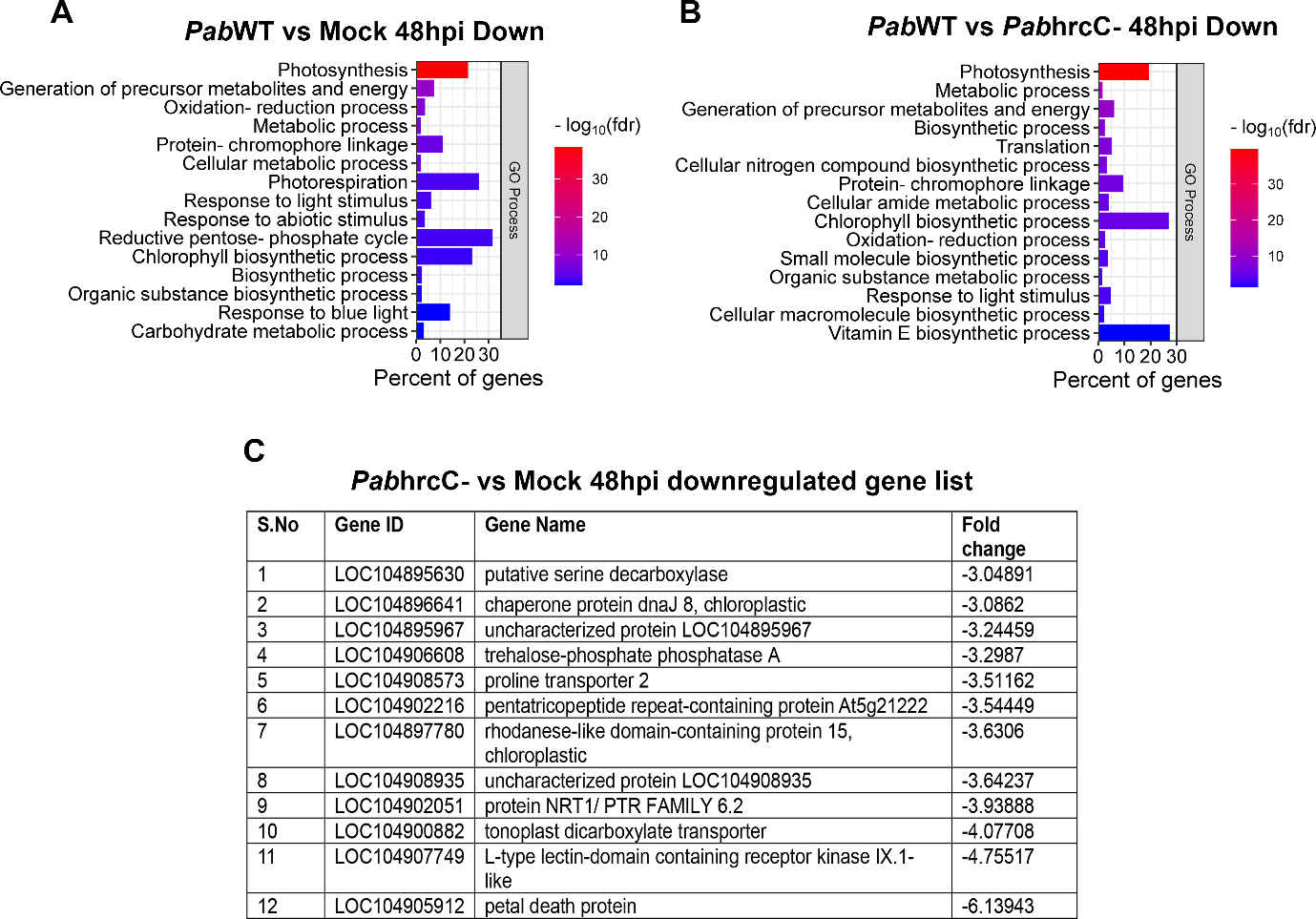

**Fig. S1 Gene ontology (GO) process enrichment analysis of downregulated genes after *Pab*WT vs mock; *Pab*hrcC- vs mock and *Pab*WT vs *Pab*hrcC^-^ treatment at 48 hpi in beet leaves.** GO functional categories enriched for downregulated genes after (A) *Pab*WT vs mock 48 hpi (B) *Pab*WT vs *Pab*hrcC^-^ treatment 48 hpi (C) Table for DEGs after *Pab*hrcC^-^ vs mock comparison at 48 hpi (FDR < 0.05, fold change ≥ 3).
