## Supplementary material for "Comparative transcriptomic profile reveals candidate genes manipulated by type III effectors of *Pantoea agglomerans* pv. *betae* leading to gall formation in beet": Fig. S2

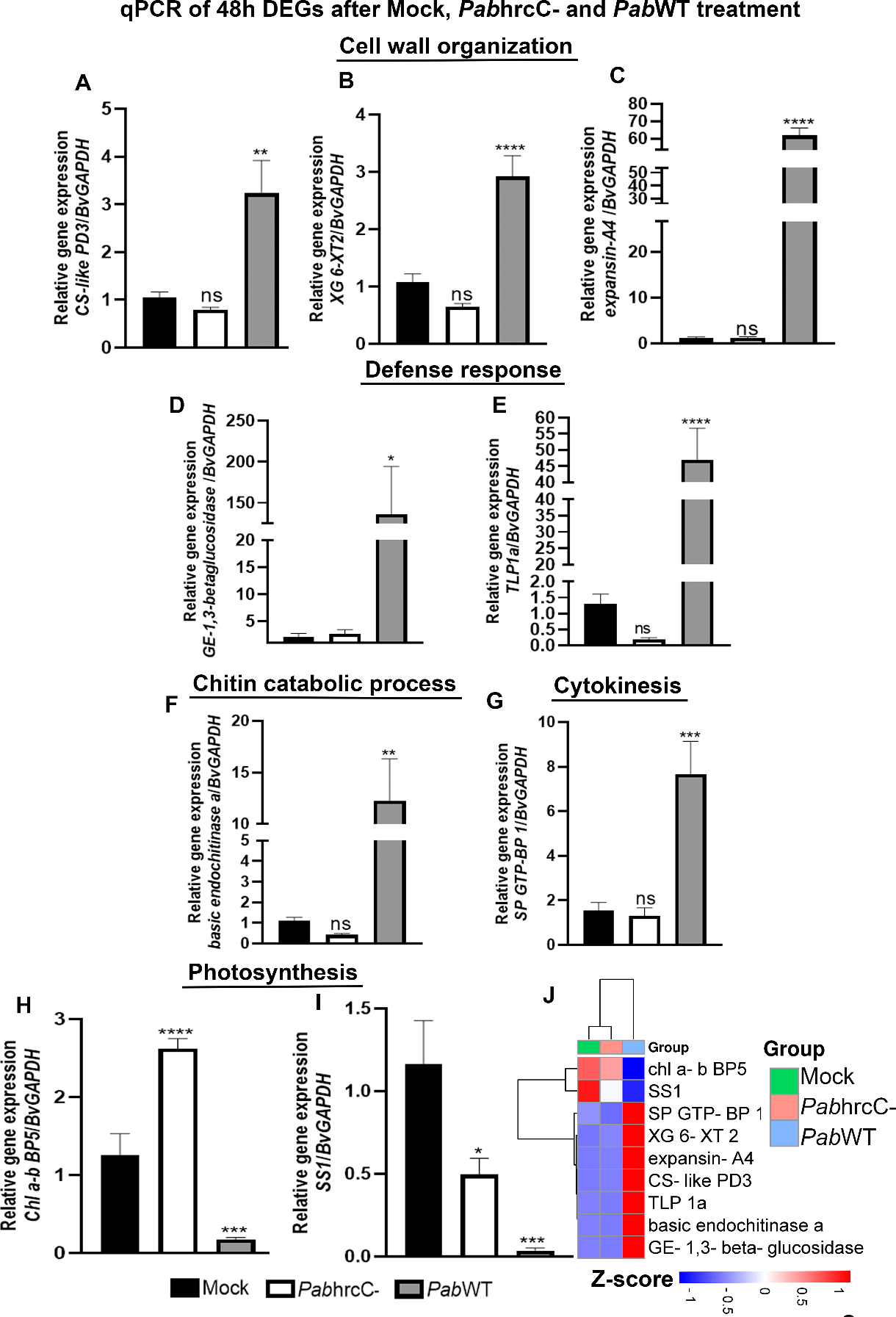

**Fig. S2**. **RT-qPCR for RNA sequencing validation of selected DEGs after 48 hpi.** (A-I) Representative genes from different functional categories were taken for validation of RNA sequencing results using RT-qPCR. (J) Heatmap for FPKM (average of replicates) values of the selected genes for validation. Asterisks represents statistically significant difference of the treatments compared to mock based on three biological replicates per treatment. It is determined by one-way ANOVA method (Tukey's multiple comparisons test); ns- not significant * (p ≤ 0.05); **(p ≤ 0.01); ***(p ≤ 0.001); ****(p ≤ 0.0001). The experiments were repeated twice with similar results. Representative genes used in the analysis: *CS-like PD3*- *cellulose synthase-like protein D3* (LOC104888008); *XG 6-XT 2*- *xyloglucan 6-xylosyltransferase 2* (LOC104883721); *expansin-A4* (LOC104904256); *GE-1,3-beta-glucosidase*- *glucan endo-1,3-beta-glucosidase* (LOC104905663); *TLP1a*- *thaumatin-like protein 1* (LOC104892311); *basic endochitinase a* (LOC104894011); *SP GTP-BP 1*- *septum-promoting GTP-binding protein 1* (LOC104907043); *chl a-b BP5*- *chlorophyll a-b binding protein 5* (LOC104883168); *SS1*- *granule-bound starch synthase 1* (LOC104892645); *BvGAPDH*- *Beta vulgaris subsp. vulgaris* *glyceraldehyde-3-phosphate dehydrogenase* (LOC104893518).
