## Supplementary material for "Comparative transcriptomic profile reveals candidate genes manipulated by type III effectors of *Pantoea agglomerans* pv. *betae* leading to gall formation in beet": Table S15

**Supplementary Table 15. Quality assessment of raw RNA-seq data**

| Sample | Total Raw Reads (Mb) | Total Clean Reads (Mb) | Total Clean Bases (Gb) | Clean reads  Q20 (%) | Clean reads  Q30 (%) | Clean reads Ratio (%) |
| --- | --- | --- | --- | --- | --- | --- |
| Mock_12h_1 | 50.83 | 42.97 | 6.45 | 97.61 | 90.27 | 84.54 |
| Mock_12h_2 | 49.08 | 42.07 | 6.31 | 97.56 | 90.08 | 85.73 |
| Mock_12h_3 | 50.83 | 42.15 | 6.32 | 97.67 | 90.39 | 82.91 |
| Mock_48h_2 | 49.08 | 43.02 | 6.45 | 97.79 | 90.75 | 87.65 |
| Mock_48h_3 | 49.08 | 42.08 | 6.31 | 97.69 | 90.48 | 85.73 |
| Mock_48h_4 | 50.83 | 43.16 | 6.47 | 96.94 | 88.38 | 84.91 |
| *Pab*WT_12h_1 | 50.83 | 42.77 | 6.42 | 97.6 | 90.22 | 84.13 |
| *Pab*WT_12h_2 | 50.83 | 42.79 | 6.42 | 97.61 | 90.22 | 84.18 |
| *Pab*WT_12h_3 | 47.33 | 42.08 | 6.31 | 97.68 | 90.43 | 88.91 |
| *Pab*WT_48h_2 | 49.08 | 42.43 | 6.36 | 96.72 | 87.74 | 86.45 |
| *Pab*WT_48h_3 | 49.08 | 42.92 | 6.44 | 96.83 | 87.95 | 87.45 |
| *Pab*WT_48h_4 | 49.08 | 43.37 | 6.5 | 96.79 | 87.95 | 88.36 |
| *Pab*hrcC_12h_1 | 50.83 | 42.44 | 6.37 | 97.66 | 90.41 | 83.49 |
| *Pab*hrcC_12h_2 | 50.83 | 43.19 | 6.48 | 97.64 | 90.32 | 84.97 |
| *Pab*hrcC_12h_3 | 49.08 | 42.24 | 6.34 | 97.68 | 90.37 | 86.06 |
| *Pab*hrcC_48_2 | 50.83 | 42.62 | 6.39 | 97.15 | 89 | 83.84 |
| *Pab*hrcC_48_3 | 49.08 | 42.21 | 6.33 | 96.78 | 87.95 | 85.99 |
| *Pab*hrcC_48_4 | 49.08 | 43.33 | 6.5 | 96.14 | 86.08 | 88.28 |
