## Supplementary material for "Comparative transcriptomic profile reveals candidate genes manipulated by type III effectors of *Pantoea agglomerans* pv. *betae* leading to gall formation in beet": Table S16

**Supplementary Table 16**. **Primers used in this study**.

| **S.No** | **Gene Name** | **Locus ID** | **Primers** |
| --- | --- | --- | --- |
| 1 | *septum-promoting GTP-binding protein 1* | (LOC104907043) | F-TGCCAAACTGGCAAAACTAGC  R- AAAGCAATGCGAGCACCTTT |
| 2 | *xyloglucan 6-xylosyltransferase 2* | (LOC104883721) | F- CGACCTGTTTTCGAGGCTGA  R- CCGAATCCAGGGCGGTATTT |
| 3 | *cellulose synthase-like protein D3* | (LOC104888008) | F- CTGGTGCTCTCACAATCGGT  R- CCCCATTCGGTCTTGTCCTC |
| 4 | *4,5-DOPA dioxygenase extradiol 1* | (LOC104908371) | F- AGCACATCCTTTCCCAGAACA  R- TGTAGGAGCCGTGACACAAG |
| 5 | *thaumatin-like protein 1* | (LOC104892311) | F- CCCATGTCCTTCAGACCCAA  R- CGGTGGTCTTCCACCTTTGT |
| 6 | *basic endochitinase* | (LOC104894011) | F- CGTGCGAATTCGATCCTTCG  R- TGGGACCCAACAAAGCCATT |
| 7 | *chlorophyll a-b binding protein 5,* | (LOC104883168) | F- AGAATGAGCTCGTCCGCAAG  R- GGGTCTGCTCTGAGAAGGGA |
| 8 | *granule-bound starch synthase 1* | (LOC104892645) | F- AGGCTGGGATCTTGGAGTCT  R- CAGAGTAGCCACACGACAGG |
| 9 | *expansin-A4* | (LOC104904256) | F- GTACAGCCAAGGGTATGGGG  R- GGAAGGACTTCCTGGGTGAC |
| 10 | *glucan endo-1,3-beta-glucosidase* | (LOC104905663) | F- ATCAGAAGGTGGGAACGCTG  R- CCAAAATGCTGCTCAGTGGC |
| 11 | *glutathione S-transferase* | (LOC104883042) | F – TGCCTCCCCTTCATTACGTG  R- TTCTCCCAAGAAGGCCTAGC |
| 12 | *peroxidase 4-like* | (LOC104908359) | F- GAGCAAGTAACTCGGGGGAC  R- TTTGGACGAGGGCATTGGTG |
| 13 | *dirigent protein 22* | (LOC104890203) | F- TACCGGTTAGTGGTGCAACG  R- ACCCAACAATGAGAATCCCTGT |
| 14 | *F-box/kelch-repeat protein At2g44130* | (LOC104889095) | F- ATTTTGGGTCGTAAGCGGGT  R- CGGGTTGATACCCGGATGAG |
| 15 | *JINGUBANG* | (LOC104897218) | F- ATTCTCGGGAGCTTGTGACC  R- AGCATAGAATCGCCGCATCA |
| 16 | *auxin-binding protein ABP19a* | (LOC104887796) | F- TGCAAGAACATCGAAATCAAGC  R-AGCATGAGATATAGACACCAAGAG |
| 17 | *glyceraldehyde-3-phosphate dehydrogenase* | (LOC104893518) | F- GCTTTGAACGACCACTTCGC  R- ACGCCGAGAGCAACTTGAAC |
